# Semantic scene-object consistency modulates N300/400 EEG components, but does not automatically facilitate object representations

**DOI:** 10.1101/2021.08.19.456466

**Authors:** Lixiang Chen, Radoslaw Martin Cichy, Daniel Kaiser

**Author notes:** R.M.C. and D.K. contributed equally to this study. **Correspondence:** Lixiang Chen, Department of Education and Psychology, Freie Universität Berlin, Berlin, Germany.

## Abstract

During natural vision, objects rarely appear in isolation, but often within a semantically related scene context. Previous studies reported that semantic consistency between objects and scenes facilitates object perception, and that scene-object consistency is reflected in changes in the N300 and N400 components in EEG recordings. Here, we investigate whether these N300/400 differences are indicative of changes in the cortical representation of objects. In two experiments, we recorded EEG signals while participants viewed semantically consistent or inconsistent objects within a scene; in Experiment 1, these objects were task-irrelevant, while in Experiment 2, they were directly relevant for behavior. In both experiments, we found reliable and comparable N300/400 differences between consistent and inconsistent scene-object combinations. To probe the quality of object representations, we performed multivariate classification analyses, in which we decoded the category of the objects contained in the scene. In Experiment 1, in which the objects were not task-relevant, object category could be decoded from around 100 ms after the object presentation, but no difference in decoding performance was found between consistent and inconsistent objects. By contrast, when the objects were task-relevant in Experiment 2, we found enhanced decoding of semantically consistent, compared to semantically inconsistent, objects. These results show that differences in N300/400 components related to scene-object consistency do not index changes in cortical object representations, but rather reflect a generic marker of semantic violations. Further, our findings suggest that facilitatory effects between objects and scenes are task-dependent rather than automatic.

## Introduction

In the real world, objects rarely appear in isolation, but practically always within a particular scene context (Bar 2004; Wolfe et al. 2011; Kaiser et al. 2019; Võ et al. 2019). Objects are often semantically related to the scene they appear in: For instance, microwaves usually appear in the kitchen, but practically never in the bathroom. Several behavioral studies have shown that such semantic relations between objects and scenes affect object identification. Early studies using line drawings of scenes and objects found that objects were detected faster and more accurately when they were in a consistent setting than in an inconsistent setting (Palmer 1975; Biederman et al. 1982; Boyce et al. 1989; Boyce and Pollatsek 1992). Similar results were recently reported for scene photographs (Davenport and Potter 2004; Davenport 2007; Munneke et al. 2013). In line with such findings, eye-tracking studies have shown that inconsistent objects are fixated longer and more often than consistent objects (Võ and Henderson 2009, 2011; Cornelissen and Võ 2017), suggesting that objects are perceived more swiftly within a consistent than within an inconsistent scene. Interestingly, such behavioral facilitation effects are also observed when, instead of the object, the scene is task-relevant: Davenport et al. (2004; 2007) reported that scenes were identified more accurately if they contained a consistent foreground object compared to an inconsistent one. These effects suggest that objects and scenes are processed in a highly interactive manner.

To characterize the neural basis of these semantic consistency effects, EEG studies have used paradigms in which objects appear within consistent or inconsistent scenes, either simultaneously or sequentially (Ganis and Kutas 2003; Mudrik et al. 2010; Võ and Wolfe 2013; Draschkow et al. 2018; Coco et al. 2020). For example, Võ and Wolfe (2013) adopted a sequential design, in which participants first viewed a scene image, followed by a location cue, where then appeared a consistent (e.g., a computer mouse on an office table) or an inconsistent object (e.g., a soap on an office table). They found objects in an inconsistent scene evoked more negative responses than consistent objects in the N300 (around 250-350 ms) and N400 (around 350-600 ms) windows. Several other studies (Mudrik et al. 2010, 2014; Truman and Mudrik 2018) using a simultaneous design, in which the scene and object were presented simultaneously, reported similar N300 and/or N400 modulations. Critically, the earlier N300 effects are often considered to reflect differences in perceptual processing between typically and atypically positioned objects (Schendan and Maher 2009; Mudrik et al. 2010; Kumar et al. 2021). On this view, consistency-related differences in EEG waveforms arise as a consequence of differences in the visual analysis of objects and scenes, rather than due to a post-perceptual signaling of (in)consistency.

If differences in the N300 waveform indeed index changes in perceptual processing, the N300 ERP effect should be accompanied by differences in the neural representation of the objects. In this study, we put this prediction to the test. Across two experiments, we compared differences in the N300/400 EEG components to multivariate decoding of objects contained in consistent and inconsistent scenes. In both experiments, participants completed a sequential semantic consistency paradigm, in which scenes from 8 different categories were consistently or inconsistently combined with objects from 16 categories. We then examined the influence of scene-object consistency on EEG signals, both when the objects were task-irrelevant (Experiment 1) and when participants performed a recognition task on the objects (Experiment 2). In both experiments, we replicated previously reported ERP effects, with greater N300 and N400 components for inconsistent scene-object combinations, compared to consistent combinations. To probe the quality of object and scene representations, we performed multivariate classification analyses, in which we decoded between the object and scene categories separately for each condition. In Experiment 1, in which the objects were not task-relevant, object category could be decoded from around 100 ms after the object presentation, but no difference in decoding performance was found between consistent and inconsistent objects. In Experiment 2, in which the objects were directly task-relevant, we found enhanced decoding of semantically consistent, compared to semantically inconsistent, objects. In both experiments, we found no differences in scene category decoding between semantically consistent and inconsistent conditions. Together, these results show that differences in N300/400 components related to scene-object consistency do not necessarily index changes in cortical object representations, but rather reflect a generic marker of semantic violations. Further, they suggest that facilitation effects between objects and scenes are task-dependent rather than automatic.

## Materials and Methods

All materials and methods were identical for the two experiments, unless stated otherwise.

### Participants

Thirty-two participants (16 males, mean age 26.23 yrs, *SD* = 2.05 yrs), with normal or corrected-to-normal vision, took part in Experiment 1. Another thirty-two participants (14 males, mean age 26.97 yrs, *SD* = 1.67 yrs) took part in Experiment 2. Participants were paid volunteers or participated for partial course credits. All participants provided written, informed consent prior to participating in the experiment. The experiments were approved by the ethical committee of the Department of Education and Psychology at Freie Universität Berlin and were conducted in accordance with the Declaration of Helsinki.

### Stimuli

The stimulus set comprised scene images from 8 categories: beach, bathroom, office, kitchen, gym, street, supermarket, and prairie. The scenes were grouped into 4 pairs (beach & bathroom, office & kitchen, gym & street, supermarket & prairie). We chose four objects for each scene pair, two of which were semantically consistent with one scene and two of which were semantically consistent with the other scene. To create semantically inconsistent scene-object combinations, we simply exchanged the objects between the scenes. All combinations of scenes and objects can be found in Table 1. For example, consider the office and kitchen pair: We first chose a computer and a printer as consistent objects for the office, and chose a rice cooker and a microwave as consistent objects for the kitchen. We in turn chose the rice cooker and microwave as inconsistent objects for the office, and the computer and printer as inconsistent objects for the kitchen. We pasted the objects into the scene images using Adobe Photoshop. The object locations were the same across the consistent and inconsistent objects, and they were always in line with the typical position of the consistent object (e.g., a computer was positioned on an office desk in the same way as a rice cooker). We used 3 exemplars for each scene category and 3 exemplars for each object, yielding 288 unique stimuli. During the experiments, the scenes could also be shown without objects (see below). Fig. 1A shows some examples of the stimuli.

**Table 1.**
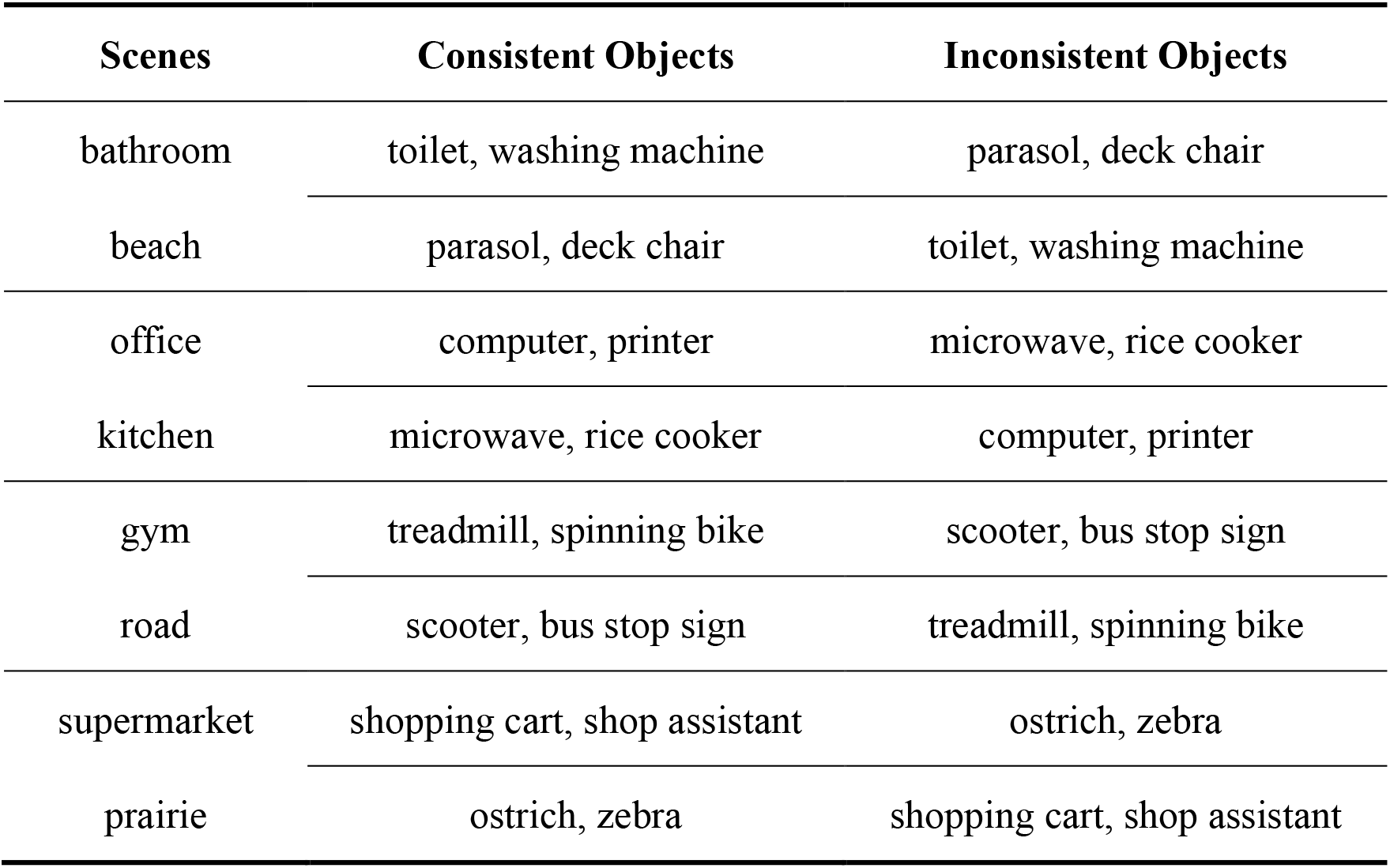
Combinations of scenes and objects for the consistent and inconsistent conditions. Note that scenes were grouped into pairs; for each pair, four objects were used as consistent and inconsistent objects.

| Scenes | Consistent Objects | Inconsistent Objects |
| --- | --- | --- |
| bathroom | toilet, washing machine | parasol, deck chair |
| beach | parasol, deck chair | toilet, washing machine |
| office | computer, printer | microwave, rice cooker |
| kitchen | microwave, rice cooker | computer, printer |
| gym | treadmill, spinning bike | scooter, bus stop sign |
| road | scooter, bus stop sign | treadmill, spinning bike |
| supermarket | shopping cart, shop assistant | ostrich, zebra |
| prairie | ostrich, zebra | shopping cart, shop assistant |

**Fig. 1.**
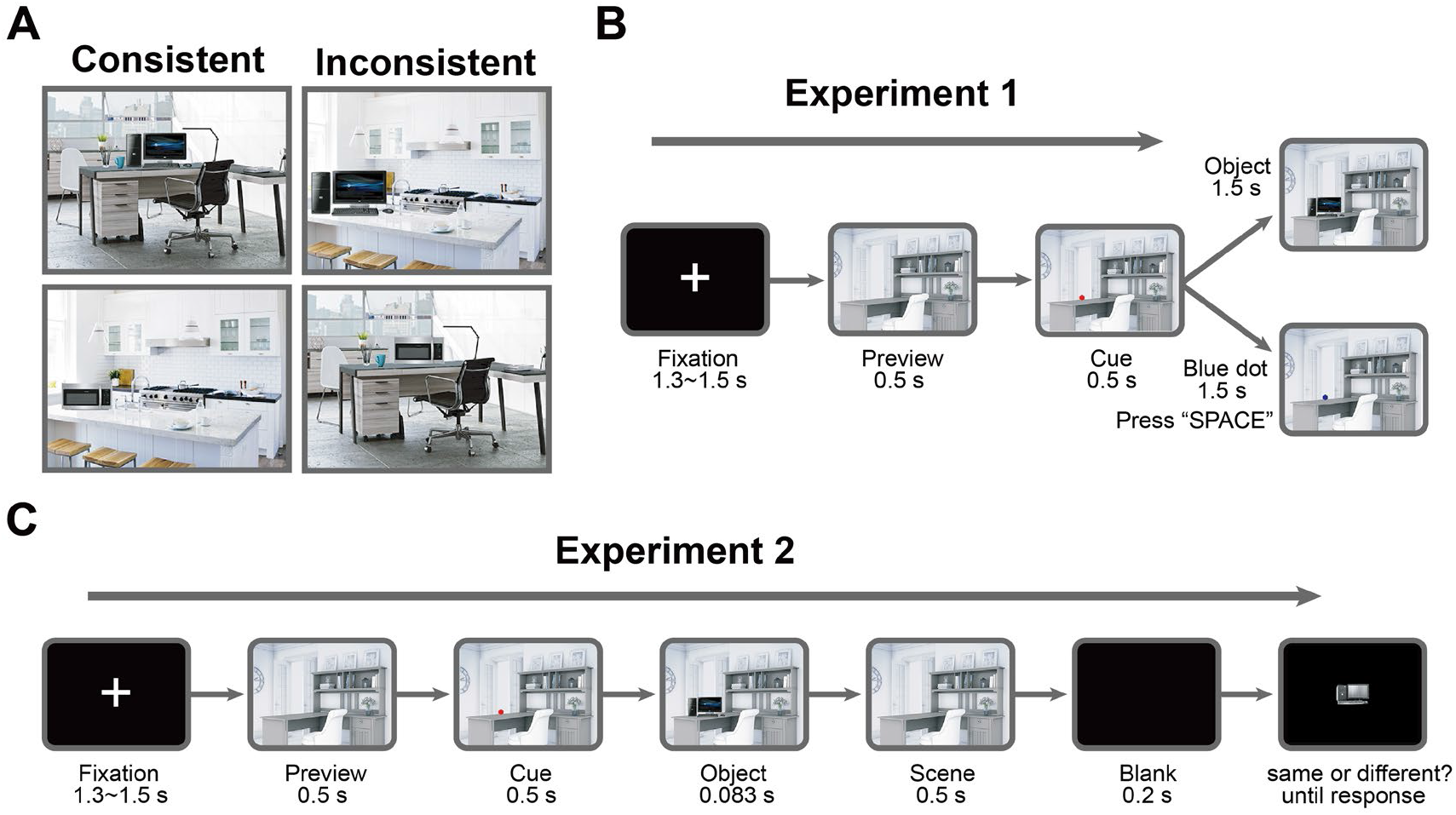
Experimental design. (A) Examples of consistent and inconsistent scene-object combinations. (B) Trial sequence in Experiment 1. After a fixation interval, a scene without the critical object was presented. Next, a red dot cue was presented, and participants were asked to move their eyes to this location. After that, the critical object appeared at the cued location in the scene. On target trials, the red cue turned blue, and participants were instructed to press spacebar. (C) Trial sequence in Experiment 2. Here, the objects were displayed briefly at the location of the cue. In a subsequent recognition test, an object of the same category appeared on the screen. Participants were asked to determine whether this object exemplar was the one that appeared earlier in the scene.

### Paradigm

Participants were seated 53 cm from an LCD monitor with a size of 34 × 27 cm and a refresh rate of 60 Hz. The images were presented on the screen at a visual angle: horizontal 20°, vertical 15.6°. We adopted a sequential scene-object congruity paradigm similar to Võ and Wolfe (2013). Image presentation and recording of subjects’ behavioral responses were controlled using MATLAB and the Psychophysics Toolbox (Brainard 1997). Each trial began with a fixation cross “+” shown for a random interval between 1.3-1.5 s, after which a scene image (without the critical object) was presented for 500 ms. Next, a red dot cue was presented at a single location in the scene for 500 ms, indicating where the critical object would appear. Participants were instructed to move their eyes to the dot as quickly as possible. To avoid eye movement artifacts during the subsequent object presentation, we told the participants not to move their eyes away from the dot location until the beginning of the next trial. After that, a semantically consistent or inconsistent object appeared at the location of the red dot.

In Experiment 1, the objects were not task-relevant. The object simply remained visible together with the scene for 1,500 ms before the next trial started. Participants performed an unrelated attention control task: To ensure that they attended the cued location, we added task trials (10% of trials) during which no object appeared after the cue. Instead, the color of the dot changed from red to blue. Participants were instructed to press the spacebar when they detected the change (detection rate in target trials: 97.8%, SE = 0.63%). An example trial for Experiment 1 is shown in Fig. 1B.

In Experiment 2, the objects were directly task-relevant. The consistent or inconsistent object appeared only very briefly (83 ms) at the location of the red dot. The scene image remained on the screen for another 500 ms after the object disappeared. After a 200 ms blank internal, participants were asked to perform an object recognition test. During the test, an object was shown on the screen, which was either the same object exemplar they had just seen or a different exemplar of the same category. Test objects were presented in grayscale to increase task difficulty. Participants were asked to determine whether this object exemplar was the one that appeared earlier in the scene. If it was, the participants should press G button, otherwise press H button. The next trial began as soon as participants made a choice. An example trial for Experiment 2 is shown in Fig. 1C.

Both experiments included three runs and all 288 unique stimulus images were presented once in random order within each run. Across runs, there were 27 repetitions for each specific scene-object combination.

### EEG Recording and Preprocessing

EEG signals were recorded using an EASYCAP 64-electrode system and a Brainvision actiCHamp amplifier in both Experiments. Electrodes were arranged according to the 10-10 system. EEG data were recorded with a sample rate of 1000 Hz and filtered online between 0.03 Hz and 100 Hz. All electrodes were referenced online to the Fz electrode and re-referenced offline to the average of data from all channels. Offline data preprocessing was performed using FieldTrip (Oostenveld et al. 2011). EEG data were segmented into epochs from −100 ms to 800 ms relative to the onset of the critical object and baseline corrected by subtracting the mean signal prior to the object onset. To track the temporal representations of scenes, EEG data were segmented into epochs from - 1100 ms to 800 ms relative to the onset of the object and baseline corrected by subtracting the mean signal prior to the scene presentation (−100-0 ms relative to the scene onset). Channels and trials containing excessive noise were removed by visual inspection. On average, we removed 1.50 ± 0.51 channels in Experiment 1 and 1.53 ± 0.57 channels in Experiment 2. These channels were not interpolated in further ERP and decoding analyses. Blinks and eye movement artifacts were removed using independent component analysis and visual inspection of the resulting components. The epoched data were downsampled to 200 Hz. As filtering is often recommended for univariate analyses but discouraged for multivariate analyses (Grootswagers et al. 2017; van Driel et al. 2021), the EEG data were not filtered for the decoding analyses. For the ERP analyses, the preprocessed data were additionally band-pass filtered at 0.1-30 Hz. This additional filtering was performed after the other preprocessing steps to equate the preprocessing pipeline between the ERP and decoding analyses. Performing the filtering before epoching yielded highly similar ERP results (see Supplementary Fig. S6 and S7).

### ERP Analyses

To replicate semantic consistency ERP effect reported in previous scene studies (e.g., Võ and Wolfe 2013; Mudrik et al. 2014), we performed ERP analyses using FieldTrip. In accordance with Võ and Wolfe (2013), we chose 9 electrodes (FC1, FCz, FC2, C1, Cz, C2, CP1, CPz, and CP2) located in the mid-central region for further ERP analysis. This a-priori electrode selection was corroborated in a topographical analysis of the scene-consistency effect (see Supplementary Fig. S5). We first averaged the evoked responses across these electrodes and then averaged these mean responses separately for the consistent and inconsistent conditions and each participant.

### Decoding Analyses

We performed two complementary multivariate decoding analyses to track temporal representations of objects and scenes across time. First, to track representations of objects and investigate how consistent or inconsistent scene contexts affect objects processing, we performed decoding analyses between two consistent and inconsistent objects separately within each scene at each time point from −100 ms to 800 ms relative to the onset of the object. For example, we performed classification analyses to either differentiate printers (consistent) from computers (consistent) in office scenes, or to differentiate printers (inconsistent) from computers (inconsistent) in kitchen scenes, at each time point. Second, to track the impact of consistent or inconsistent objects on scenes representations, we performed decoding analyses to discriminate between every two scene categories separately for consistent and inconsistent conditions at each time point from −100 ms to 1800 ms relative to the onset of the scene (−1100 ms to 800 ms relative to the onset of the object). For example, we performed classification analyses to differentiate office scenes containing a printer or computer (consistent) from kitchen scenes containing a microwave or rice cooker (consistent), or to differentiate office scenes containing a microwave or rice cooker (inconsistent) from kitchen scenes containing a printer or computer (inconsistent). In both analyses, we used all available trials, including those in participants responded incorrectly. For each decoding analysis, we adopted two approaches: standard timeseries decoding (Boring et al. 2020; Kaiser and Nyga 2020), using data from a sliding time window, and cumulative decoding (Ramkumar et al. 2013; Kaiser, et al. 2020a), using aggregated data from all elapsed time points. The two approaches are detailed in the following paragraphs.

#### Timeseries decoding

Timeseries decoding analyses were performed using Matlab and CoSMoMVPA (Oosterhof et al. 2016). To increase the power of our timeseries decoding, the analysis was performed on a sliding time window (50 ms width), with a 5 ms resolution. This approach thus not only utilizes data from current time point, but the data from 5 time points before and after the current time point. For each sliding window position, we concatenated the response patterns across all time points within the time window and then unfolded the whole pattern into a vector. For a comparison with alternative timeseries decoding approach, which use individual time point or average data across the sliding windows, see Supplementary Fig. S2 and S3.

Considering excessive data dimensionality may harm classification, we adopted principal component analysis (PCA) to reduce the dimensionality of the data (Grootswagers et al. 2017; Kaiser and Nyga 2020; Kaiser, et al. 2020a). For each classification, a PCA was performed on all data from the training set, and the PCA solution was projected onto data from the testing set (Experiment 1, mean 34.8 PCs for object decoding, mean 52.1 PCs for scene decoding; Experiment 2, mean 41.2 PCs for object decoding, mean 69.9 PCs for scene decoding). For each PCA, we retained the set of components explaining 99% of the variance in the training set data.

The classification was performed separately for each time point from −100 to 800 ms (from −1100 to 800 ms for scene decoding), using LDA classifiers with 10-fold cross-validation. Specifically, the EEG data from all epochs were first allocated to 10 folds randomly. LDA classifiers were then trained on data from 9 folds and then tested on data from the left-out fold. The amount of data in the training set was always balanced across conditions. For each object decoding analysis, the training set included up to 48 trials, and the testing set included up to 6 trials; for each scene decoding analysis, the training set included up to 96 trials and the testing set included up to 12 trials. The classification was done repeatedly until every fold was left out once. For each time point, the accuracies were averaged across the 10 repetitions.

#### Cumulative decoding

We also performed cumulative decoding analyses, which takes into account the data of all time points before the current time point in the epoch for classifications (Ramkumar et al. 2013; Kaiser, et al. 2020a). For example, for the first time point in the epoch, the classifier was trained and tested on response patterns at this time point in the epoch; at the second time point in the epoch, the classifier was trained and tested on response patterns at the first and second time points in the epoch; and at the last time point in the epoch, the classifier was trained and tested on response patterns at all time points in the epoch. For each time point, we concatenated the response patterns across all time points up to the current one and then unfolded the whole pattern into a vector that was subsequently used for decoding.

The cumulative decoding approach uses increasingly large amounts of data that span multiple time points. This allows classifiers to capitalize on complex spatiotemporal response patterns that emerge across the trial, which may provide additional sensitivity for detecting effects that are not only visible across electrode space but that are also transported by variations in the time domain. On the flip side, this renders the interpretation of the results less straightforward: One can only conclude that spatiotemporal response patterns up to the current point allow for classification, but not which features enable this classification.

As for the timeseries decoding, LDA classifiers with 10-fold cross-validation were used for classifications and PCA was adopted to reduce the dimensionality of the data for each classification step across time (Experiment 1, mean 39.2 PCs for object decoding, mean 73.5 PCs for scene decoding; Experiment 2, mean 43.9 PCs for object decoding, mean 86.9 PCs for scene decoding).

### Statistics

For the behavioral responses in Experiment 2, we used paired *t*-tests to compare participants’ accuracy and response times when they were asked to recognize consistent and inconsistent objects.

For ERP analyses, we used paired *t*-tests to compare the averaged EEG responses evoked by consistent and inconsistent scene-object combinations, at each time point.

For decoding analyses, we used one-sample *t*-tests to compare decoding accuracies against chance level (similar results were obtained using a permutation-based testing approach, see Supplementary Fig. S1), and paired *t*-tests to compare decoding accuracies between the consistent and inconsistent conditions, at each time point.

Differences in ERP and decoding effects between experiments were assessed using independent t-tests. Direct differences between the ERP and decoding effects obtained in the two experiments were assessed in a mixed-effects ANOVA with the factors consistency effect (ERP versus decoding) and experiment (Experiment 1 versus Experiment 2).

Multiple-comparison corrections were performed using FDR (*p* < 0.05), and only clusters of at least 5 consecutive significant time points (i.e., 25 ms) were considered. We also calculated Bayes factors (Rouder et al. 2009) for all analyses.

## Results

### Experiment 1

#### ERP signals indexing scene-object consistency

To track the influence of scene-object consistency on EEG responses, we first analyzed EEG waveforms in mid-central electrodes. In this analysis, we found more negative responses evoked by inconsistent scene-object combinations than consistent combinations, which emerged at 170-330 ms (peak: *t*(31) = 4.884, BF_10_ = 765.26) and 355-470 ms (peak: *t*(31) = 3.429, BF_10_ = 20.06) (Fig. 2). These results demonstrate larger N300 and N400 components evoked by inconsistent scenes, which is in line with previous findings (Mudrik et al. 2010, 2014; Võ and Wolfe 2013).

**Fig. 2.**
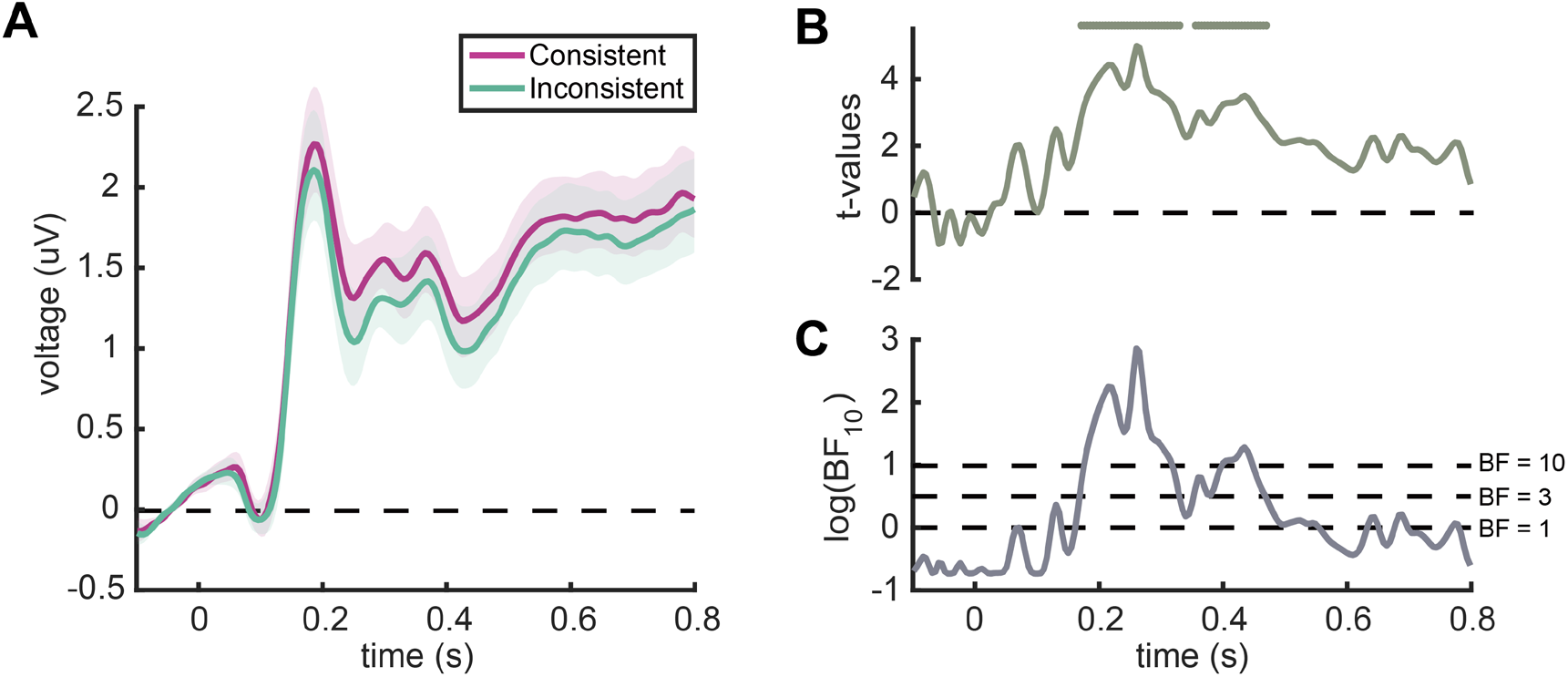
Event-related potentials (ERPs) in Experiment 1. **(A)** ERPs recorded from the mid-central region for consistent and inconsistent scene-object combinations. Error margins represent standard errors. **(B)** *t*-values for the comparisons between consistent and inconsistent conditions. Line markers denote significant differences between conditions (*p* < 0.05, FDR-corrected). **(C)** Bayes factors (BF_10_) for the comparisons between consistent and inconsistent conditions. For display purposes, the BF_10_ values were log-transformed. Dotted lines show low (BF_10_ = 1), moderate (BF_10_ = 3), and high (BF_10_ = 10) evidence for a difference between conditions. In line with previous reports, these results show that scene-object consistency is represented in evoked responses 170-330 ms and 355-470 ms after the object onset.

#### Tracking object representations in consistent and inconsistent scenes

Having established reliable ERP differences between consistent and inconsistent scene-object combinations, we were next interested if these differences were accompanied by differences in how well the consistent and inconsistent objects were represented. We performed timeseries and cumulative decoding analyses between two consistent or inconsistent objects separately within each scene at each time point from −100 to 800 ms relative to the onset of the object. In both analyses, we found highly similar decoding performances for both consistent and inconsistent objects. Specifically, there was significant decoding between consistent objects, which emerged at 65-790 ms in the timeseries decoding (Fig. 3A), and between 80 and 800 ms in the cumulative decoding (Fig. 3D), and there was significant decoding between inconsistent objects in both the timeseries decoding (60-645 ms; Fig. 3A) and cumulative decoding (90-800 ms; Fig. 3D). No significant differences in decoding accuracy were found between consistent and inconsistent objects. Hence, despite the reliable ERP differences between consistent and inconsistent scene-object stimuli, there was no evidence for an automatic facilitation from the scene to the semantically consistent object.

**Fig. 3.**
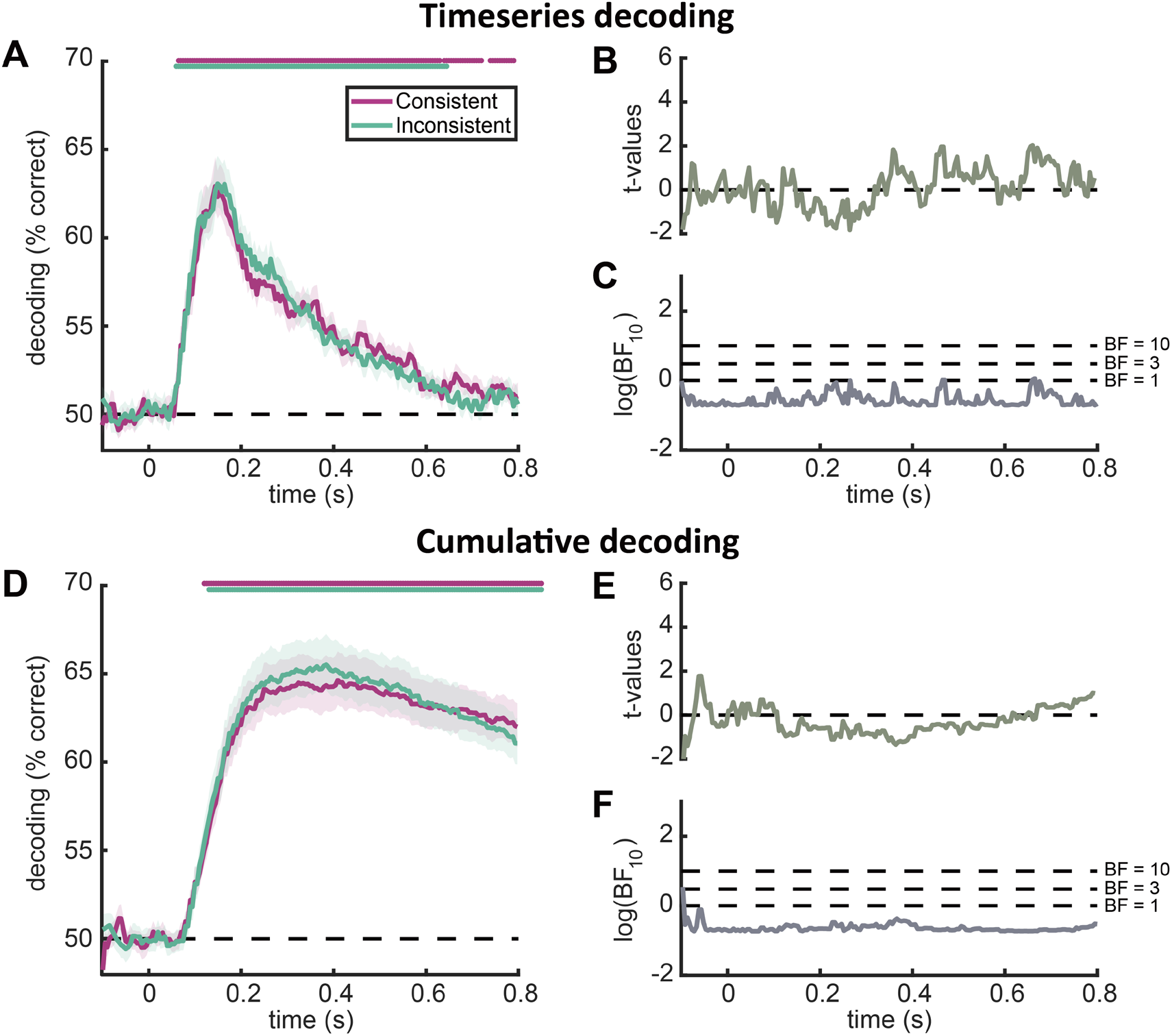
Decoding for the consistent or inconsistent objects within each scene in Experiment 1. **(A)** Timeseries decoding results, separately for consistent and inconsistent objects. Line markers denote significant above-chance decoding (*p* < 0.05, FDR-corrected). **(B)** *t*-values for the comparisons between consistent and inconsistent conditions. **(C)** Bayes factors (BF_10_) for the comparisons between consistent and inconsistent conditions. For display purposes, the BF_10_ values were log-transformed. Dotted lines show low (BF_10_ = 1), moderate (BF_10_ = 3), and high (BF_10_ = 10) evidence for a difference between conditions. **(D)** Cumulative decoding results, separately for consistent and inconsistent objects. Line markers denote significant above-chance decoding (*p* < 0.05, FDR-corrected). **(E, F)** *t*-values and Bayes factors (BF_10_) for the comparisons between consistent and inconsistent conditions, as in (B, C). These results show robust decoding for consistently and inconsistently placed objects. However, despite the reliable differences between consistent and inconsistent scene-object combinations in ERP signals, object decoding was highly similar between the consistent and inconsistent conditions.

#### Tracking the representation of scenes with consistent and inconsistent objects

Although scene-object consistency does not automatically facilitate the representation of the objects, there may still be an opposite cross-facilitation effect where the consistent object enhances scene representations. To test this possibility, we performed decoding analyses to discriminate between every two categories separately for the consistent and inconsistent conditions from −100 ms to 1800 ms relative to the onset of the scene. We found significant decoding between scenes with a consistent object in both the timeseries decoding (45-1800 ms) and cumulative decoding (80-1800 ms) analyses. Significant decoding between scenes that contained inconsistent objects was also found in both the timeseries decoding (40-1800 ms) and cumulative decoding (80-1800 ms). These results are consistent with previous findings (Lowe et al. 2018; Kaiser, et al. 2020b), which suggest scene category can be decoded within 100 ms (Fig. 4). However, no significant differences were found between these scenes with consistent and inconsistent objects. Such differences were also not observed when we corrected for multiple comparisons solely between 0-800 ms relative to the onset of the object. These results suggest that scene category can be decoded in a temporally sustained way, but semantically consistent objects have no facilitatory effect on scene representations.

**Fig. 4.**
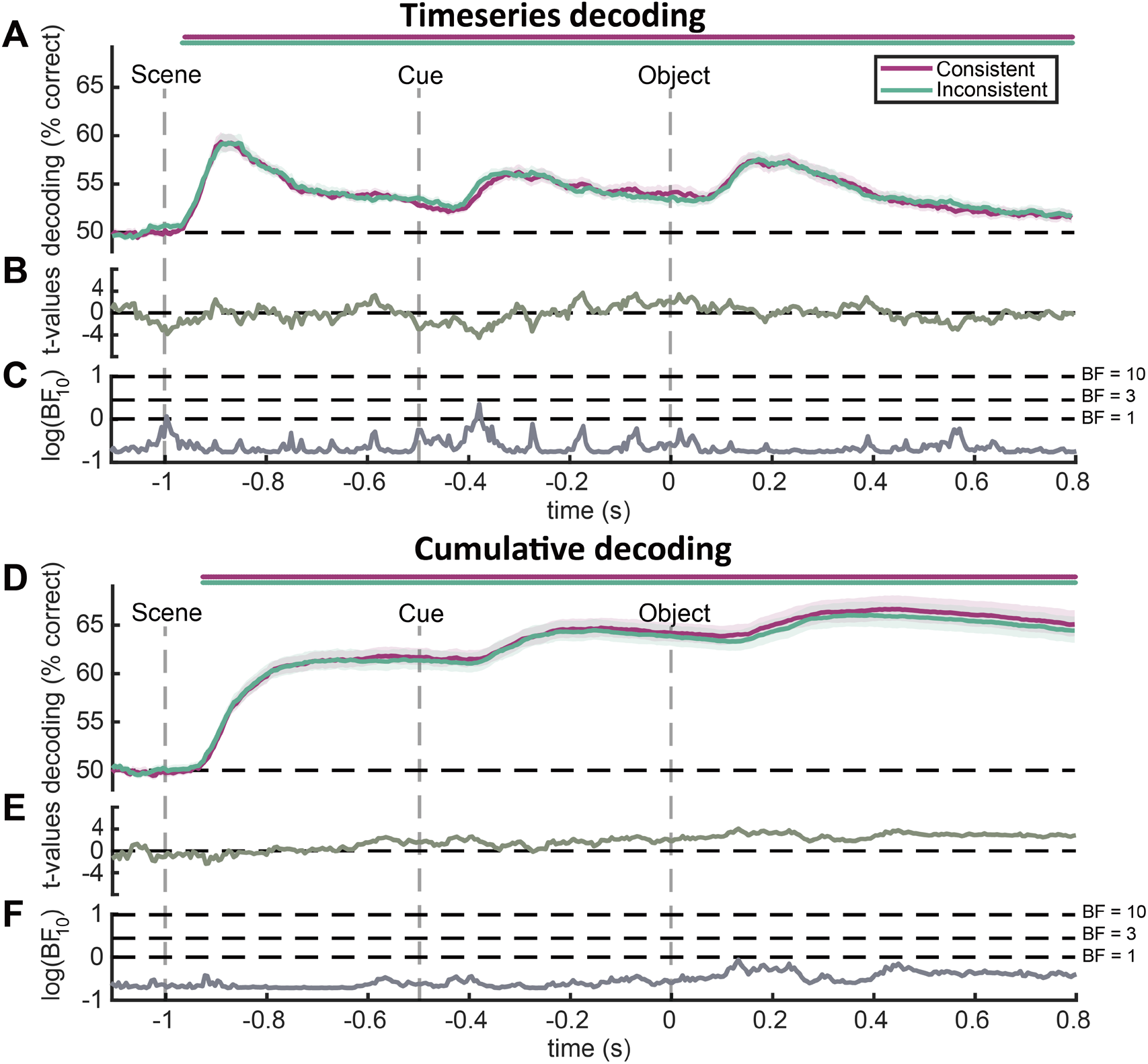
Decoding between scenes with consistent or inconsistent objects in Experiment 1. **(A)** Timeseries decoding results, separately for scene with consistent and inconsistent objects. Line markers denote significant above-chance decoding (*p* < 0.05, FDR-corrected). **(B)** *t*-values for the comparisons between consistent and inconsistent conditions. **(C)** Bayes factors (BF_10_) for the comparisons between consistent and inconsistent conditions. For display purposes, the BF_10_ values were log-transformed. Dotted lines show low (BF_10_ = 1), moderate (BF_10_ = 3), and high (BF_10_ = 10) evidence for a difference between conditions. **(D)** Cumulative decoding results, separately for scene with consistent and inconsistent objects. Line markers denote significant above-chance decoding (*p* < 0.05, FDR-corrected). **(E, F)** *t*-values and Bayes factors (BF_10_) for the comparisons between consistent and inconsistent conditions, as in (B, C). These results show that scene category can be well decoded across time, but the consistency of embedded objects has no facilitatory effects on scene representations.

### Experiment 2

In Experiment 1, we did not find differences in object and scene representations between the consistent and inconsistent object-scene combinations, despite robust ERP differences between the two conditions. However, the objects and scenes were both not task-relevant in Experiment 1 – although participants spatially attended the object location, the objects’ features were not important for solving the task. Under such conditions, object representations may not benefit from semantically consistent context to the same extent as when object features are critical for solving the task. In Experiment 2, we therefore made the objects task-relevant.

#### Behavioral object recognition in semantically consistent and inconsistent scenes

In Experiment 2, participants performed a recognition task, in which they were asked to report whether a test object was identical to the one they had previously seen in the scene (Fig. 1C). In line with previous findings (Davenport and Potter 2004; Davenport 2007; Munneke et al. 2013), we found that objects were recognized more accurately when they were embedded in consistent scenes than in inconsistent scenes (mean accuracy: consistent = 82.36%, inconsistent = 79.30%; *t*(31) = 2.598, *p* = 0.011). These results suggest that semantically consistent scenes can enhance the recognition of objects. There was no difference in response times between two conditions (mean response time: consistent = 723.8 ms, inconsistent = 742.4 ms; *t*(31) = −0.648, *p* = 0.519).

#### ERP signals indexing scene-object consistency

Inconsistent scene-object combinations evoked more negative responses in mid-central electrodes than consistent combinations at 240-335 ms (peak: *t*(31) = 4.385, BF_10_ = 210.85), 360-500 ms (peak: *t*(31) = 4.291, BF_10_ = 165.94), and 570-590 ms (peak: *t*(31) = 2.986, BF_10_ = 7.36) (Fig. 5). The results suggest larger N300 and N400 components evoked by semantically inconsistent scene-object combinations, replicating the findings from Experiment 1.

**Fig. 5.**
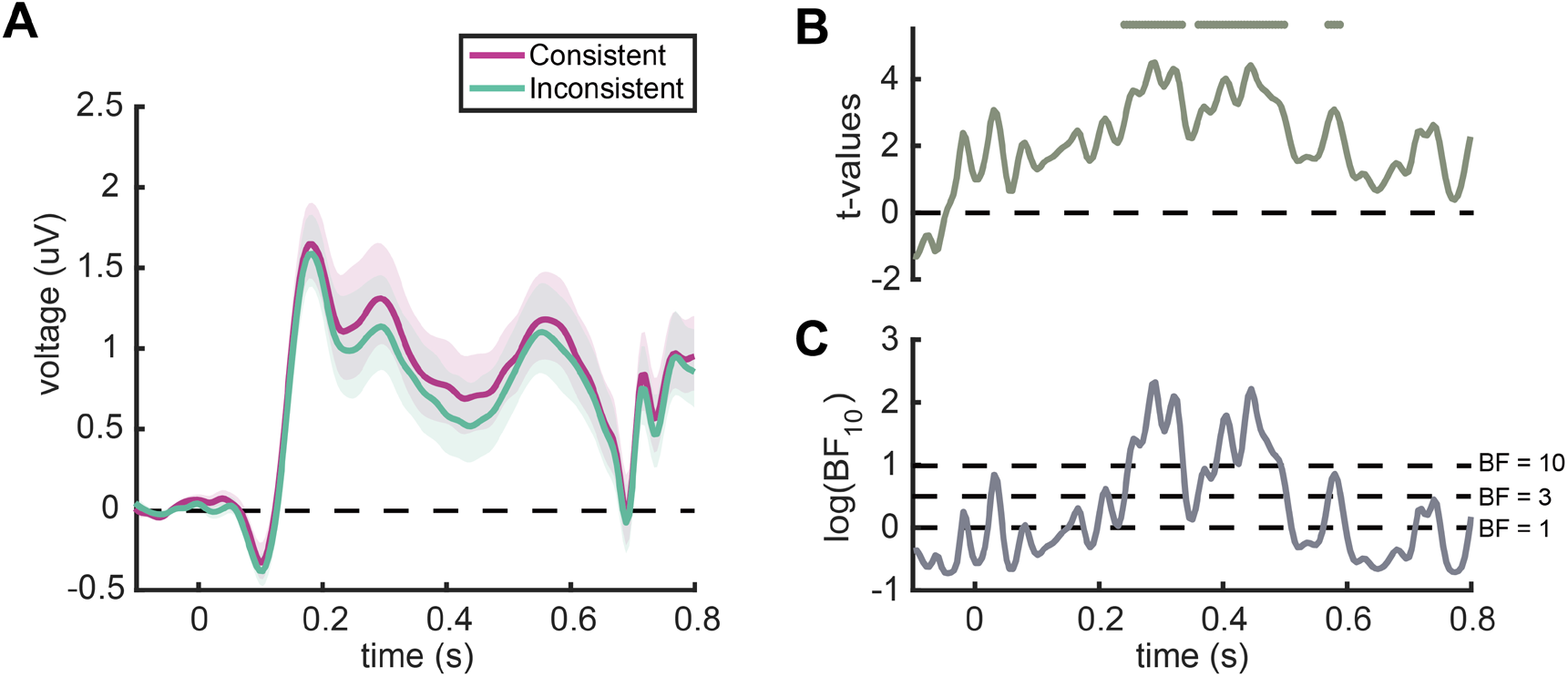
Event-related potentials (ERPs) in Experiment 2. (A) ERPs recorded from the mid-central region for consistent and inconsistent scene-object combinations. Error margins represent standard errors. (B) t-values for the comparisons between consistent and inconsistent conditions. Line markers denote significant differences between conditions (p < 0.05, FDR-corrected). (C) Bayes factors (BF10) for the comparisons between consistent and inconsistent conditions. For display purposes, the BF10 values were log-transformed. Dotted lines show low (BF10 = 1), moderate (BF10 = 3), and high (BF10 = 10) evidence for a difference between conditions. Similar to Experiment 1, inconsistent scene-object combinations evoked more negative responses at 240-335 ms, 360-500 ms, and 570-590 ms after the object onset relative to consistent combinations.

#### Tracking object representations in consistent and inconsistent scenes

To test whether semantically consistent scenes facilitate object representations differently from semantically inconsistent scenes when the objects are task-relevant, we performed both timeseries and cumulative decoding analyses, where we classified two consistent or inconsistent objects within each scene at each time point from −100 to 800 ms relative to the onset of the object. We found significant decoding for both consistent objects (timeseries decoding: 60-800 ms; cumulative decoding: 70-800 ms) and inconsistent objects (timeseries decoding: 70-760 ms; cumulative decoding: 90-800 ms). Critically, we found the consistent objects were decoded more accurately than inconsistent objects in both the timeseries decoding (310-410 ms and 545-680 ms) and cumulative decoding analyses (160-190 ms, 255-295 ms, and 370-800 ms) (Fig. 6). Additional electrode-space searchlight analyses suggest that these enhanced representations primarily emerge in posterior electrodes over the right hemisphere (see Supplementary Fig. S4). These results suggest that scene-object consistency can facilitate cortical object representations – but only when the objects are task-relevant. Our data show that such effects arise at least from around 300ms, although the more sensitive cumulative decoding suggests that such effects may be seen much earlier, even within the first 200ms of processing. As the current evidence for such early effects is only moderately strong, the exact timing of such effects needs to be confirmed in future studies.

**Fig. 6.**
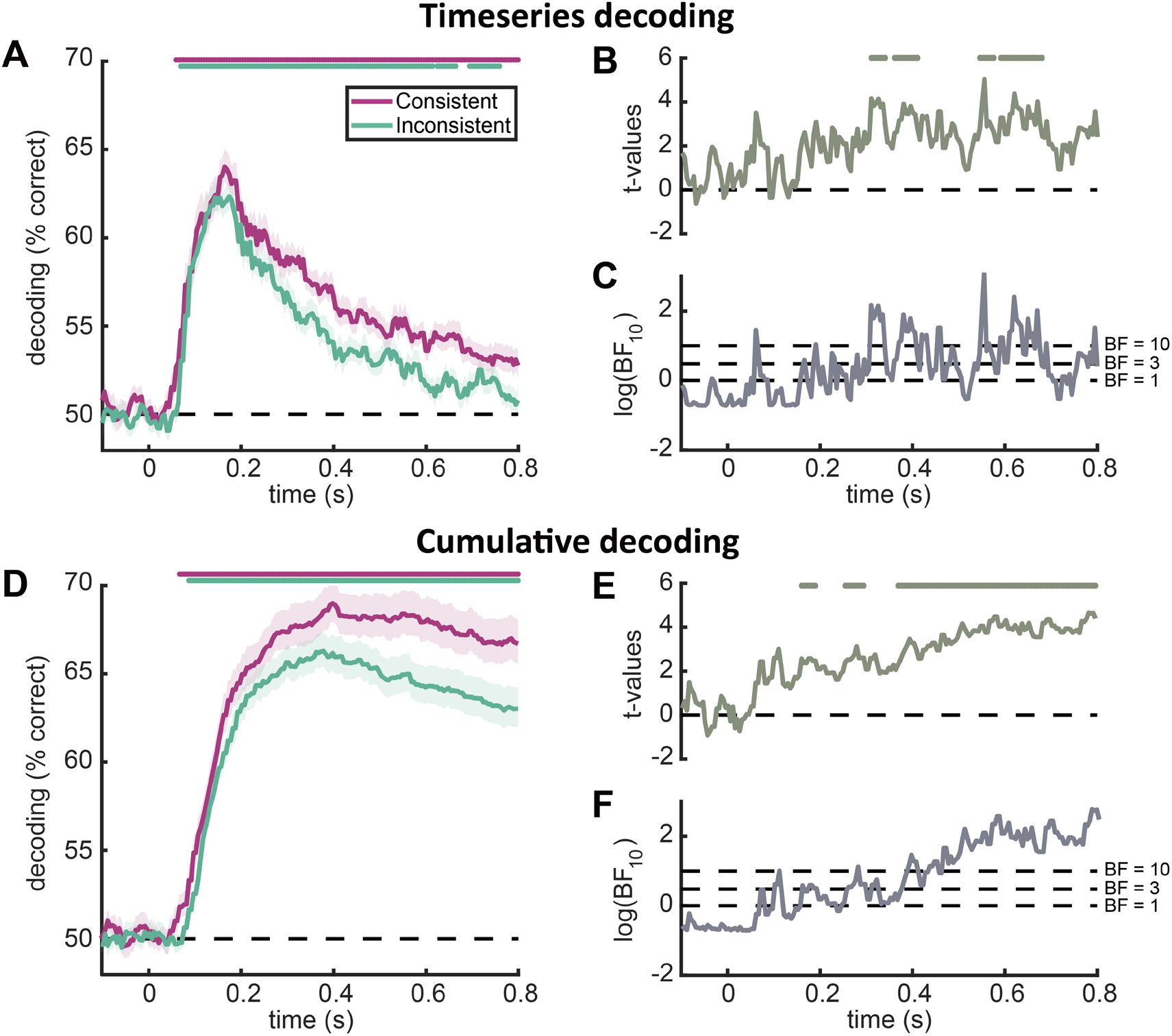
Decoding for the consistent or inconsistent objects within each scene in Experiment 2. **(A)** Timeseries decoding results, separately for consistent and inconsistent objects. Line markers denote significant above-chance decoding (*p* < 0.05, FDR-corrected). **(B)** *t*-values for the comparisons between consistent and inconsistent conditions. Line markers denote significant differences between the consistent and inconsistent conditions (*p* < 0.05, FDR-corrected). **(C)** Bayes factors (BF_10_) for the comparisons between consistent and inconsistent conditions. For display purposes, the BF_10_ values were log-transformed. Dotted lines show low (BF_10_ = 1), moderate (BF_10_ = 3), and high (BF_10_ = 10) evidence for a difference between conditions. **(D)** Cumulative decoding results, separately for consistent and inconsistent objects. Line markers denote significant above-chance decoding (*p* < 0.05, FDR-corrected). **(E, F)** *t*-values and Bayes factors (BF_10_) for the comparisons between consistent and inconsistent conditions, as in (B, C). These results are markedly different from Experiment 1: Scenes can indeed facilitate the cortical representations of consistent objects when the objects are task-relevant.

#### Tracking the representation of scenes with consistent and inconsistent objects

As in Experiment 1, we also tested whether semantically consistent objects can facilitate scene representations. We performed timeseries and cumulative decoding analyses to discriminate between every two scene categories separately for the consistent and inconsistent conditions from −100 ms to 1800 ms relative to the onset of the scene. We found very similar results as the Experiment 1, with significant decoding for both consistent scenes (timeseries decoding: 55-1800 ms; cumulative decoding: 85-1800 ms) and inconsistent scenes (timeseries decoding: 60-1800 ms; cumulative decoding: 85-1800 ms), but no difference in decoding performance between consistent and inconsistent conditions (Fig. 7). As in Experiment 1, such differences were also not observed when we corrected for multiple comparisons solely between 0-800 ms relative to the onset of the object. These results suggest that facilitation effects between scenes and objects are not mutual, but that they likely depend on behavioral goals: once the objects were task relevant, we found a facilitation effect originating from semantically consistent scenes.

**Fig. 7.**
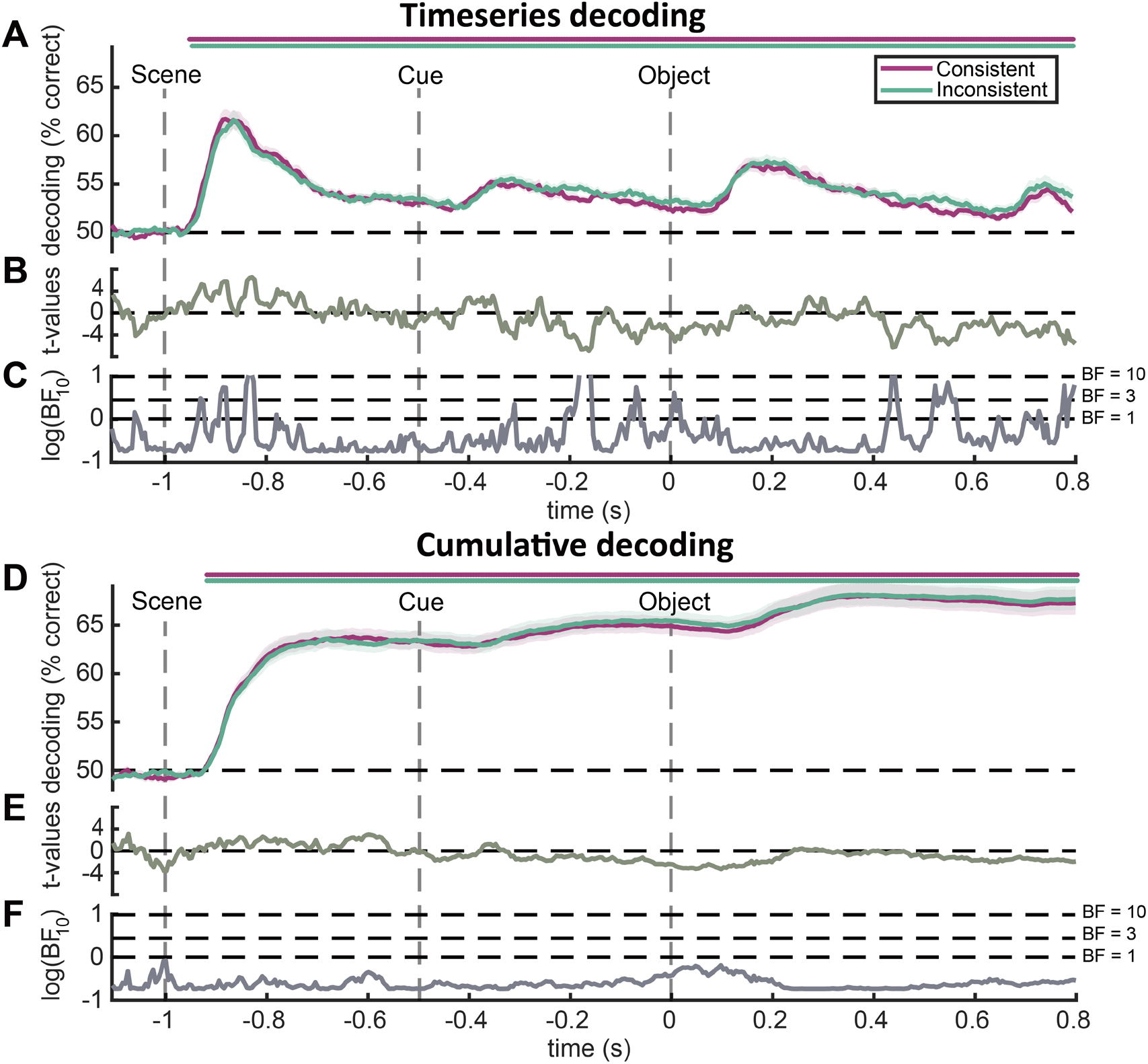
Decoding between scenes with consistent or inconsistent objects in Experiment 2. **(A)** Timeseries decoding results, separately for scene with consistent and inconsistent objects. Line markers denote significant above-chance decoding (*p* < 0.05, FDR-corrected). **(B)** *t*-values for the comparisons between consistent and inconsistent conditions. **(C)** Bayes factors (BF_10_) for the comparisons between consistent and inconsistent conditions. For display purposes, the BF_10_ values were log-transformed. Dotted lines show low (BF_10_ = 1), moderate (BF_10_ = 3), and high (BF_10_ = 10) evidence for a difference between conditions. **(D)** Cumulative decoding results, separately for scene with consistent and inconsistent objects. Line markers denote significant above-chance decoding (*p* < 0.05, FDR-corrected). **(E, F)** *t*-values and Bayes factors (BF_10_) for the comparisons between consistent and inconsistent conditions, as in (B, C). The results show that consistent embedded objects do not automatically facilitate the representation of scenes.

#### Comparison across experiments

The pattern of results across our experiments revealed reliable ERP effects that are independent of task-relevance, but multivariate decoding demonstrated that representational facilitation effects can only be observed when the objects are task relevant. To statistically quantify this pattern, we directly compared the ERP and decoding results between two experiments. For each participant, we computed the difference between the consistent and inconsistent conditions, and then compared these differences across experiments using independent-samples *t*-tests. For the ERP results, we found no statistical differences across the experiments (all *p* > 0.05, FDR-corrected; Fig. 8A), suggesting that N300/400 effects emerge independently of the task-relevance of the objects. On the flipside, object representations benefitted more strongly from semantically consistent context when the objects were directly task-relevant: In the timeseries decoding, differences between the two experiments emerged at 215-245 ms, 295-335 ms, and 610-635 ms (max *t*(62) = 3.40, *p* = 0.001), suggesting that during these time points task-relevance enhances the effect of semantically consistent scene context (Fig. 8B). Clear evidence for this effect was also found in the cumulative decoding, where the effect of semantically consistent scene context was stronger in Experiment 2 between 160-190 ms and 255-800 ms (Fig. 8C). There were no significant differences in scene decoding between the two experiments.

**Fig. 8.**
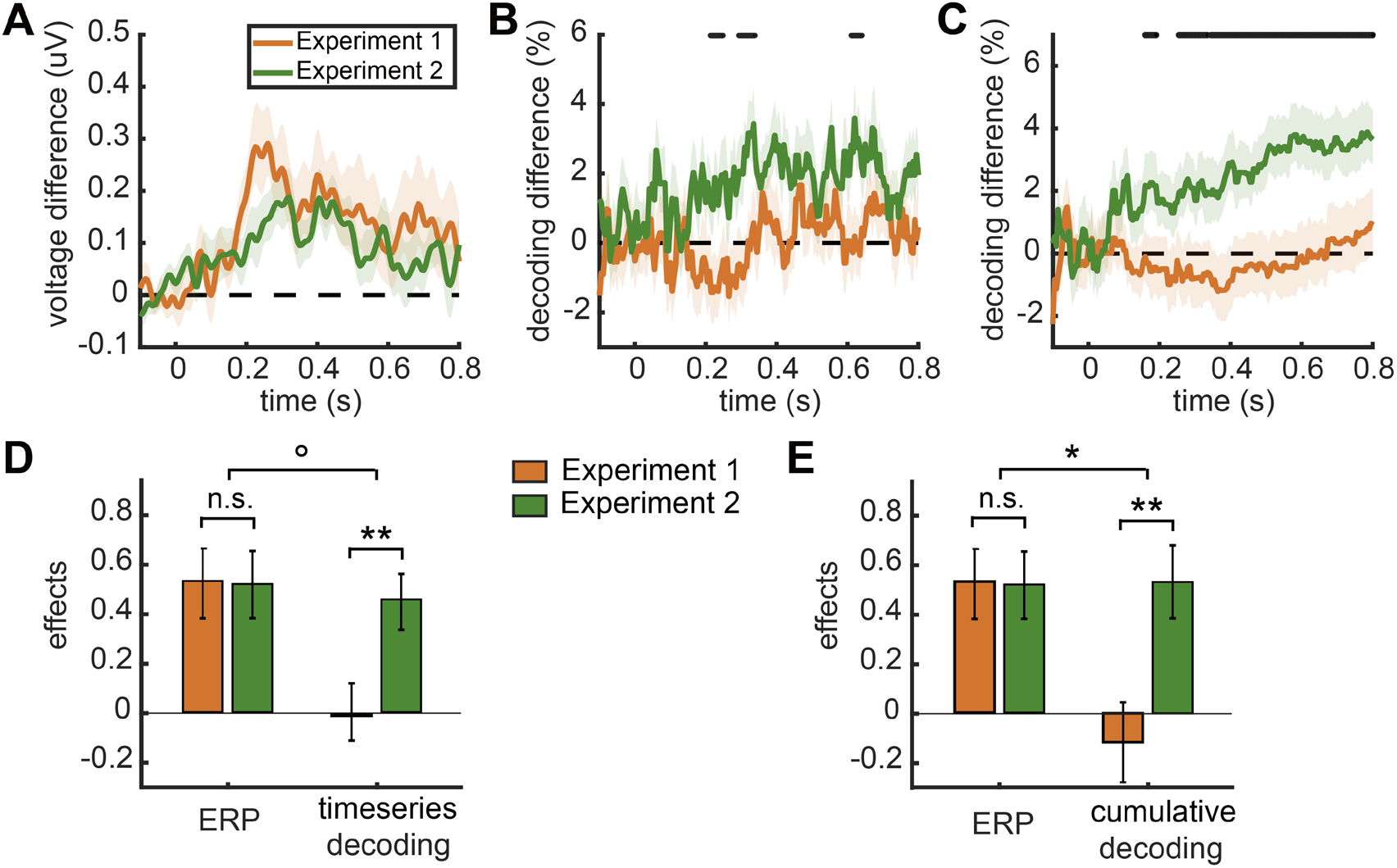
Comparisons of ERP and decoding effects between experiments. **(A)** Differences in ERP effects (consistent – inconsistent) between experiments. **(B)** Differences in decoding effects (consistent – inconsistent) for object in timeseries decoding analyses between experiments. Line markers denote significant differences between two experiments (*p* < 0.05). **(C)** Differences in decoding effects for object in cumulative decoding analyses between experiments. Line markers denote significant differences between two experiments (*p* < 0.05). These results shows that the N300/400 effects emerge independently of the task relevance of the objects, but facilitations from scenes to the representations of objects are task-dependent. **(D)** Standardized consistency effects in ERPs and timeseries object decoding in both experiments, averaged across the time span during which ERP effects were observed in the two experiments (170-590 ms). **(E)** Standardized effects in ERPs and cumulative object decoding in both experiments, averaged across 170-590 ms. The difference between ERP and decoding effects in the two experiments was assessed using a mixed effects ANOVA. Error bars represent standard error of the mean. ° represents *p* < 0.1; * represents *p* < 0.05; ** represents *p* < 0.01.

Finally, we asked whether the difference in results across experiments was statistically different for the ERP and the object decoding analyses. To answer this question, we performed a 2 × 2 mixed ANOVA with the factors task-relevance (task-irrelevant versus task-relevant objects, i.e., Experiment 1 versus Experiment 2) and neural measure (ERP versus object decoding). For this analysis, we first needed to make the ERP and decoding effects comparable. To achieve this, we first calculated the ERP difference between consistent and inconsistent conditions at each time point for each participant and standardized the difference values by dividing them by the standard deviation of ERP differences within the group at each time point. The effects for timeseries and cumulative object decoding were calculated and standardized in the same way. Next, to obtain a more reliable estimate of the ERP and decoding effects, we used the time span in which the ERP effects emerged in both experiments (between 170 ms and 590 ms after object onset) to average the ERP and decoding effects across time. Finally, we performed 2 × 2 mixed ANOVAs to test the interaction effect, separately for the timeseries and cumulative decoding results. The expected interaction between task-irrelevance/relevance and ERP/decoding failed to reach significance when looking at the timeseries decoding, but revealed a trend (*F*(1, 62) = 3.009, *p* = 0.088; Fig. 8D). When comparing between ERPs and cumulative decoding, the interaction reached significance (*F*(1, 62) = 4.296, *p* = 0.042; Fig. 8E). This result indicates the effect of task-relevance is significantly larger in the cumulative object decoding than it is in the ERP analysis. This corroborates the notion that the N300/400 effects are dissociable from changes in object representation, as indexed by our decoding analyses.

Together, this shows that N300/400 ERP differences emerge independently of task relevance, suggesting that they do not index changes in object representations. By contrast, multivariate decoding reveals that changes in object representations are modulated by task-relevance: when the objects are critical for behavior, semantically consistent scenes more strongly enhance their cortical representation.

## Discussion

In this study, we used EEG to investigate how scene-object consistency affects the quality of object and scene representations. In two experiments, we replicated previous scene-object consistency ERP effects (Mudrik et al. 2010, 2014; Võ and Wolfe 2013), showing that inconsistent scene-object combinations evoked more negative responses in the N300 and N400 windows than consistent combinations. Critically, multivariate decoding analyses revealed whether these scene-object consistency effects in ERPs were accompanied by changes in the quality of cortical object and scene representations. We found that task-irrelevant consistent and inconsistent objects were decoded equally well in Experiment 1, despite pronounced ERP differences in the N300/400 range. When the objects were task-relevant in Experiment 2, we observed a comparable N300/400 ERP effect, which now was accompanied by enhanced object decoding. Across both experiments, we found no significant differences in scene category decoding between consistent and inconsistent conditions. These results suggest that the N300/400 ERP effects are not necessarily indicative of enhanced object or scene representations. Further, they suggest that facilitations between objects and scenes are task-dependent rather than automatic.

### N300 effects do not index changes in perceptual processing

The N300 effects found in the study replicated previous findings in studies of scene-object consistency (Võ and Wolfe 2013; Mudrik et al. 2014; Draschkow et al. 2018; Truman and Mudrik 2018; Coco et al. 2020). Particularly the early N300 effects were often interpreted as reflecting differences in perceptual processing (Schendan and Maher 2009; Mudrik et al. 2010; Dyck and Brodeur 2015; Sauvé et al. 2017; Kumar et al. 2021). Such findings are often explained through models of contextual facilitation (Bar, 2004; Bar et al., 1996), which propose that object representations are refined by more readily available information about the consistent context. Specifically, when a scene is presented, gist-consistent schemas are rapidly activated through non-selective processing channels (Wolfe et al. 2011). By comparing this rapidly available scene gist to incoming visual information, perceptual uncertainty in object recognition is reduced. However, if the object does not match the scene gist, its identification should be impeded. It was argued that this mismatch between inconsistent objects and the pre-activated schemas elicits a larger N300 amplitude, signifying a prediction error that occurs during perceptual object analysis (Kumar et al. 2021).

Our data challenge this interpretation. We show that enhanced N300 amplitudes are observed independently of changes in object decoding. We found reliable N300 differences between consistent and inconsistent objects, which were highly similar for task-relevant and task-irrelevant objects. By contrast, object information, as measured by our multivariate object decoding analyses, was similar for consistent and inconsistent objects when they were not task-relevant; only when they were task-relevant, we found that scene-object consistency facilitated object representations. It is worth noting that both task-relevant and task-irrelevant objects within the scenes could be decoded reliably and with high accuracy in both experiments, which is in line with previous reports (Kaiser et al. 2016); our results therefore cannot be attributed to a failure to decode the objects in the first place.

The pattern of results obtained in our study is therefore inconsistent with the N300 indexing a change in perceptual representations. Our results are rather consistent with an interpretation that views the N300 as a general marker of inconsistency or a purely attentional response to a violation of expectation. On this view, N300 differences are post-perceptual in nature. Contrary to the N300, consistency-related differences in the N400 time window are commonly interpreted as a marker of differences in post-perceptual semantic processing (Võ and Wolfe 2013; Truman and Mudrik 2018). In fact, a recent study has shown that N400 effects are qualitatively similar to N300 effects (Draschkow et al. 2018), further supporting the view that N300 differences are not directly indicative of changes in perceptual encoding.

When interpreting the results of our study, two limitations should be considered. First, to render the object-level task sufficiently difficult, we drastically reduced the presentation time of the object in Experiment 2 (from 1,500 ms to 83 ms). There is thus a possibility that the longer timing, rather than the lack of behavioral relevance, in Experiment 1 abolished decoding effects in Experiment 1. Further experiments are needed to establish a clear distinction between these explanations. However, if the ERP effects and the decoding effects were indeed a reflection of the same underlying changes in perceptual representations, they should both be affected by object timing – given that we did not observe any change in ERP effects across the two experiments, but a marked difference in object decoding, it is unlikely that the timing difference concealed an otherwise tight correspondence between the two effects. Further, we observed highly comparable overall decoding in the two experiments, which suggests that longer presentation times do not per se alter object decoding. Second, our study compares univariate ERP analyses that average many trials on a small set of electrodes with multivariate decoding analyses that probe a set of pairwise combinations between conditions across large-scale electrode patterns. Ultimately, these different approaches yield different (unknown) sensitivities, so that a comparison between results obtained with the two approaches within a single experiment can be challenging. However, the different results obtained across our two experiments cannot be attributed to an overall sensitivity difference between methods.

### Semantic consistency only facilitates task-relevant representations

Our results suggest that cross-facilitation effects between objects and scenes are not automatic but task-dependent. Consistent objects were only decoded better than inconsistent objects in Experiment 2 where they were directly task-relevant, suggesting that semantically consistent scenes only facilitate object processing when the objects are critical for behavior. Further, decoding between the different scenes was similar for scenes that contained consistent and inconsistent objects. As the scenes were never task-relevant, this supports the view that that mutual influences between scene and object representations are only observed when they support ongoing behavior.

Several previous neuroimaging studies reported a cross-facilitation between scene and object processing (Brandman and Peelen 2017, 2019; Kaiser et al. 2021), reporting that scenes enhance the cortical representation of objects (Brandman and Peelen 2017; Kaiser et al. 2021), and objects facilitate the representation of scenes (Brandman and Peelen 2019). In these studies, participants were asked to attend the objects or scenes by memorizing them, completing repetition detection tasks, or categorization tasks. One recent study directly compared cross-facilitation effects between objects and scenes under different task demands (Kaiser et al. 2021). In this study, spatially consistent scene context facilitated object representation more than spatially inconsistent scene context when the objects were task-relevant. When participants instead performed a task on the scene, object representations were comparable for the spatially consistent and inconsistent scene contexts.

These results are in line with the current study, in which semantically consistent scene context only facilitated perceptual object processing when it was beneficial for the task at hand. Our findings therefore support a view where the visual system uses contextual information in a flexible and strategic way: When scene context is beneficial for the current task demands, the visual system harnesses contextual information to enhance object representations. Conversely, if the current task does not benefit from contextual information, no cross-facilitation between object and scene processing is found.

What mechanism governs such situational interactions between the scene and object processing systems? A recent TMS study shines new light on how contextual information from a scene may impact object processing in visual cortex (Wischnewski and Peelen 2021): By virtually lesioning key nodes of the scene and object processing networks, they established that information is first processed in parallel in object- and scene-selective cortices (until around 200 ms of processing), before information from scene-selective brain regions converges with object coding in object-selective regions (after 250 ms of processing). Our results suggest that the information flow from scene-selective to object-selective cortex is gated by behavioral demands: When the task requires perceiving object details, the brain may use scene representations to actively generate predictions about the objects that are likely to appear in the scene (Hochstein and Ahissar 2002; Bar 2004). By feeding back these predictions to object-selective cortex, the perceptual representation of consistent object information is then facilitated, for instance by sharpening the neural response (de Lange et al. 2018). When the task does not require perceiving object detail, such predictions may simply not be generated – or not generated to the same extent. This showcases how predictive processing is adaptive tailored to situational needs.

## Conclusions

In the study, we investigated how scene-object consistency affects scene and object representations. Our results suggest that differences in the N300/400 components related to scene-object consistency do not directly index differences in perceptual representations, but rather reflect a generic marker of semantic violations. Further, they suggest that facilitation effects between objects and scenes are task-dependent rather than automatic. Our findings highlight that there are multiple markers of semantic consistency that reflect different underlying brain mechanisms. How these mechanisms interact to support efficient real-world vision needs to be explored in future studies.

## Acknowledgments

The authors thank the HPC Service of ZEDAT, Freie Universität Berlin, for computing time and support.

