## Supplementary_Information for "Semantic scene-object consistency modulates N300/400 EEG components, but does not automatically facilitate object representations"

**Index of figures**

**Fig. S1** Permutation-test for timeseries object decoding against chance level.

**Fig. S2** Decoding of consistent and inconsistent objects in Experiment 1.

**Fig. S3** Decoding of consistent and inconsistent objects in Experiment 2.

**Fig. S4** Topographies of decoding accuracy for consistent and inconsistent objects in Experiments 1 and 2

**Fig. S5** Topographies of ERP differences between consistent and inconsistent scene-object combinations in Experiments 1 and 2.

**Fig. S6** Event-related potentials (ERPs) in Experiment 1, with filtering performed before epoching.

**Fig. S7** Event-related potentials (ERPs) in Experiment 2, with filtering performed before epoching.

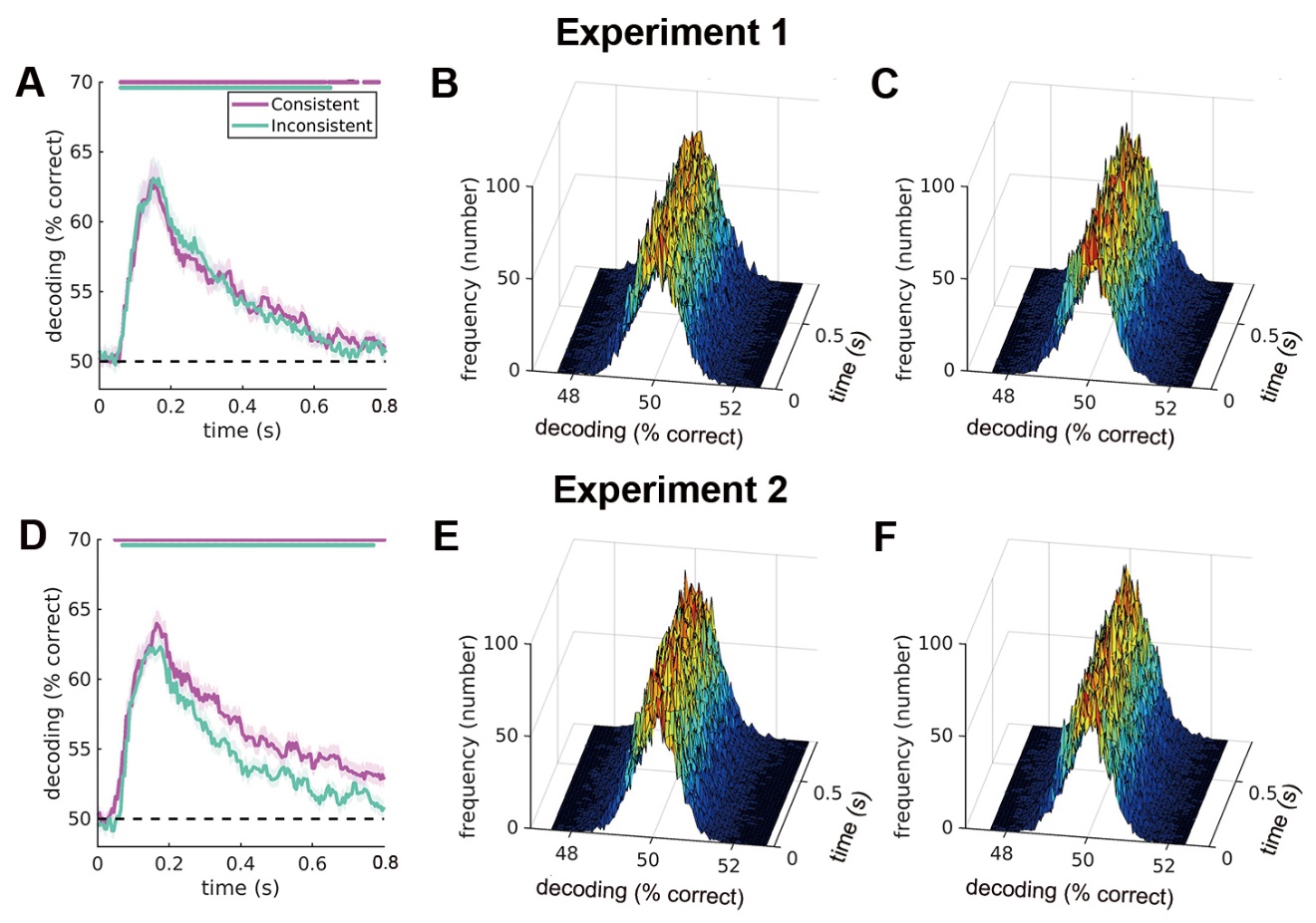

**Fig. S1** Permutation-test for timeseries object decoding against chance level. **(A)** Timeseries decoding results in Experiment 1, separately for consistent and inconsistent objects. Line markers denote significant above-chance decoding (permutation-test 1,000 times, FDR-corrected). **(B)** Null distribution for the timeseries decoding of consistent objects in Experiment 1 at each time point obtained from the permutation test. **(C)** Null distribution for the timeseries decoding of inconsistent objects in Experiment 1 at each time point obtained from the permutation test. **(D)** Timeseries decoding results in Experiment 2. **(E)** Null distribution for the timeseries decoding of consistent objects in Experiment 2 at each time point. **(F)** Null distribution for the timeseries decoding of inconsistent objects in Experiment 2 at each time point. These results show the empirical chance levels for the timeseries decoding of both consistent and inconsistent objects in both experiments are centered around 50% and the significant timepoints are virtually identical to our original approach.

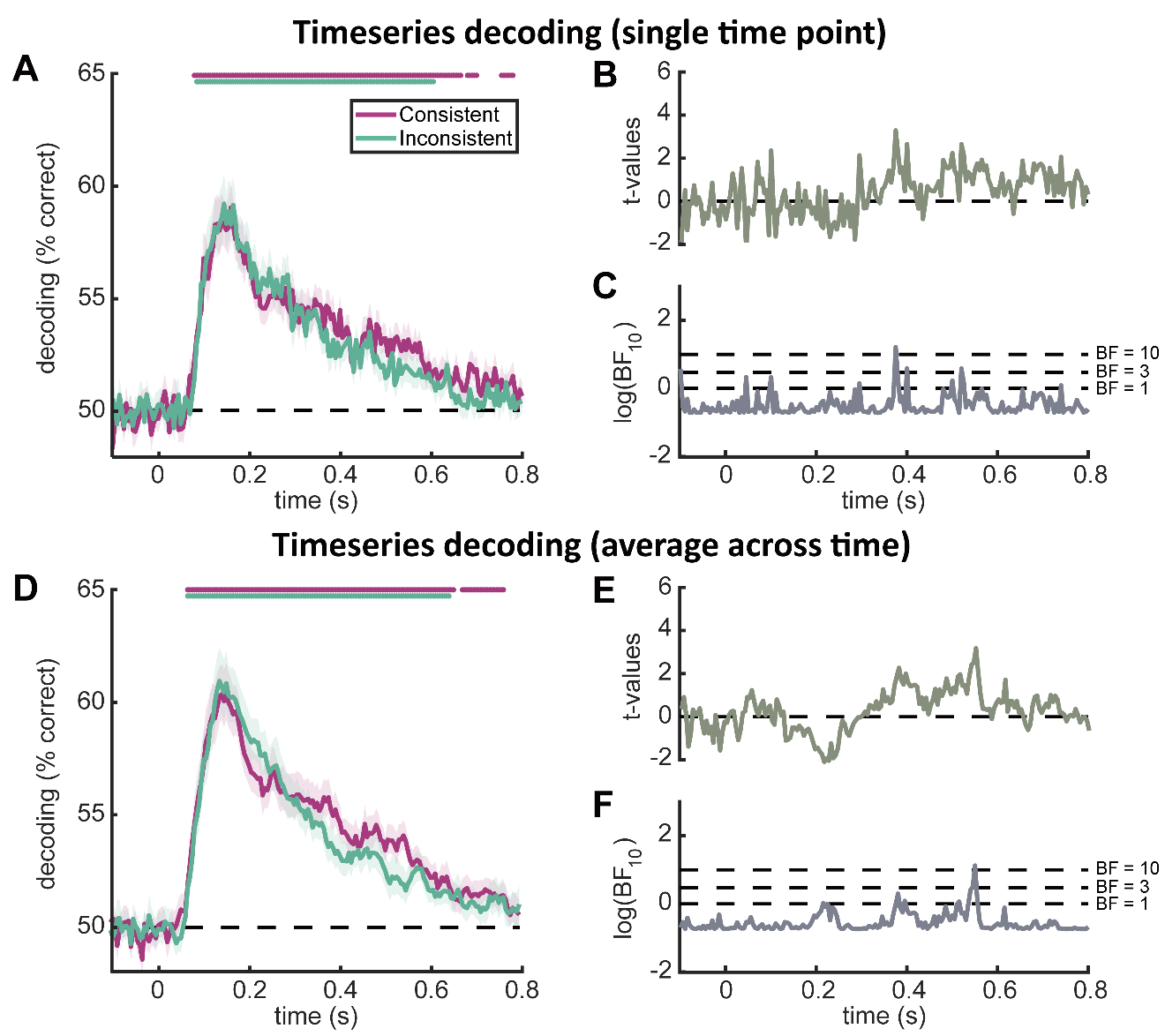

**Fig. S2** Decoding of consistent and inconsistent objects in Experiment 1. **(A)** Results of timeseries decoding using the data from a single time point without using sliding windows, separately for consistent and inconsistent objects. Line markers denote significant above-chance decoding (*p* < 0.05, FDR-corrected). **(B)** *t*-values for the comparisons between consistent and inconsistent conditions. **(C)** Bayes factors (BF_10_) for the comparisons between consistent and inconsistent conditions. For display purposes, the BF_10_ values were log-transformed. Dotted lines show low (BF_10_ = 1), moderate (BF_10_ = 3), and high (BF_10_ = 10) evidence for a difference between conditions. **(D)** Results of timeseries decoding using the data averaged across time within each 50 ms sliding window, separately for consistent and inconsistent objects. **(E, F)** *t*-values and Bayes factors (BF_10_) for the comparisons between consistent and inconsistent conditions, as in (B, C). These results are very similar to the results of our timeseries decoding using sliding windows.

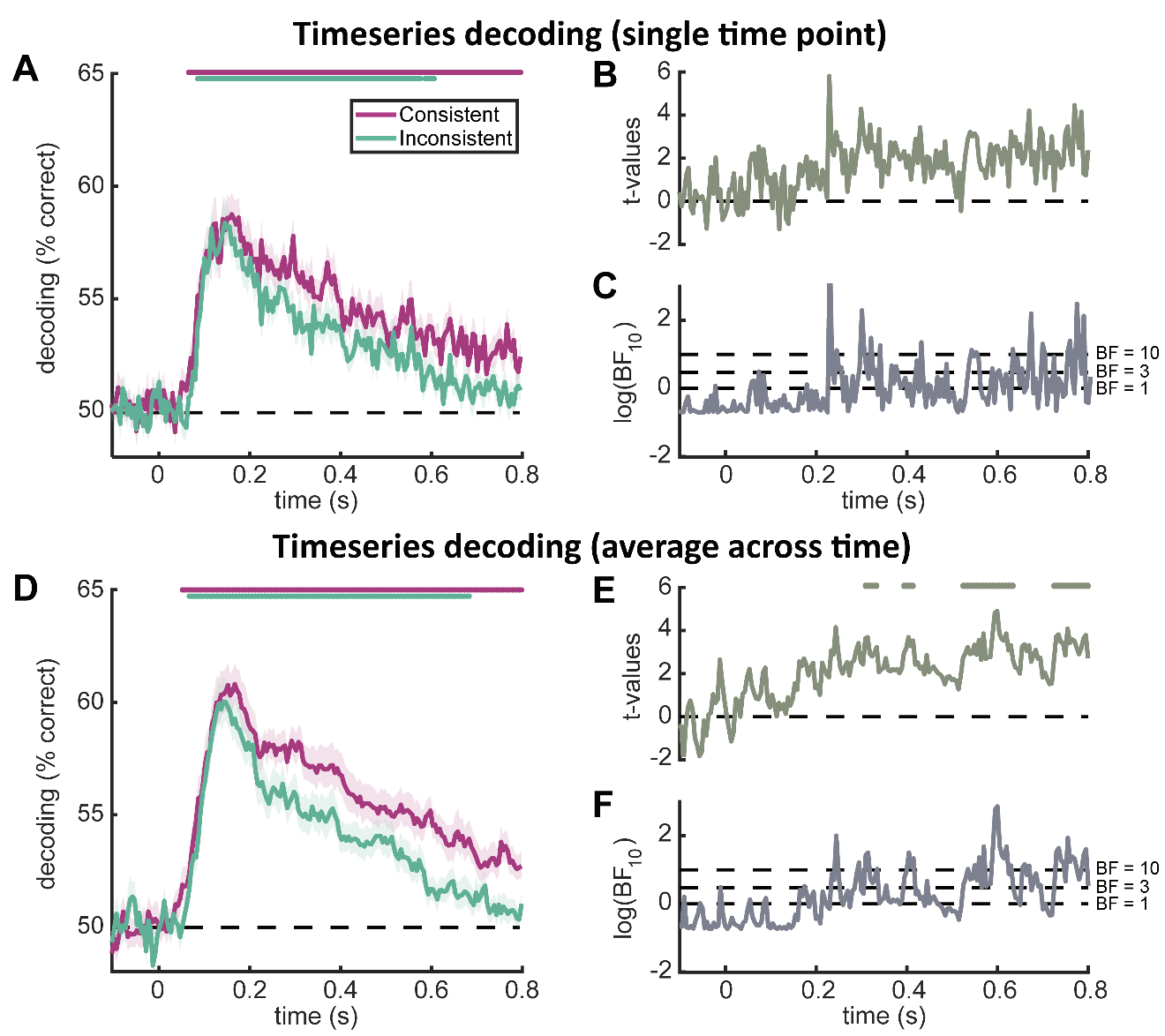

**Fig. S3** Decoding of consistent and inconsistent objects in Experiment 2. **(A)** Results of timeseries decoding using the data from a single time point without using sliding windows, separately for consistent and inconsistent objects. Line markers denote significant above-chance decoding (*p* < 0.05, FDR-corrected). **(B)** *t*-values for the comparisons between consistent and inconsistent conditions. Line markers denote significant differences between the consistent and inconsistent conditions (*p* < 0.05, FDR-corrected). **(C)** Bayes factors (BF_10_) for the comparisons between consistent and inconsistent conditions. For display purposes, the BF_10_ values were log-transformed. Dotted lines show low (BF_10_ = 1), moderate (BF_10_ = 3), and high (BF_10_ = 10) evidence for a difference between conditions. **(D)** Results of timeseries decoding using the data averaged across time within each 50 ms sliding window, separately for consistent and inconsistent objects. **(E, F)** *t*-values and Bayes factors (BF_10_) for the comparisons between consistent and inconsistent conditions, as in (B, C). Results for the analysis on averaged time windows are similar to our original results using sliding windows: the consistent objects are decoded better than inconsistent objects when the objects are task relevant. The results on individual time points are also qualitatively similar, but the differences failed to reach statistical significance.

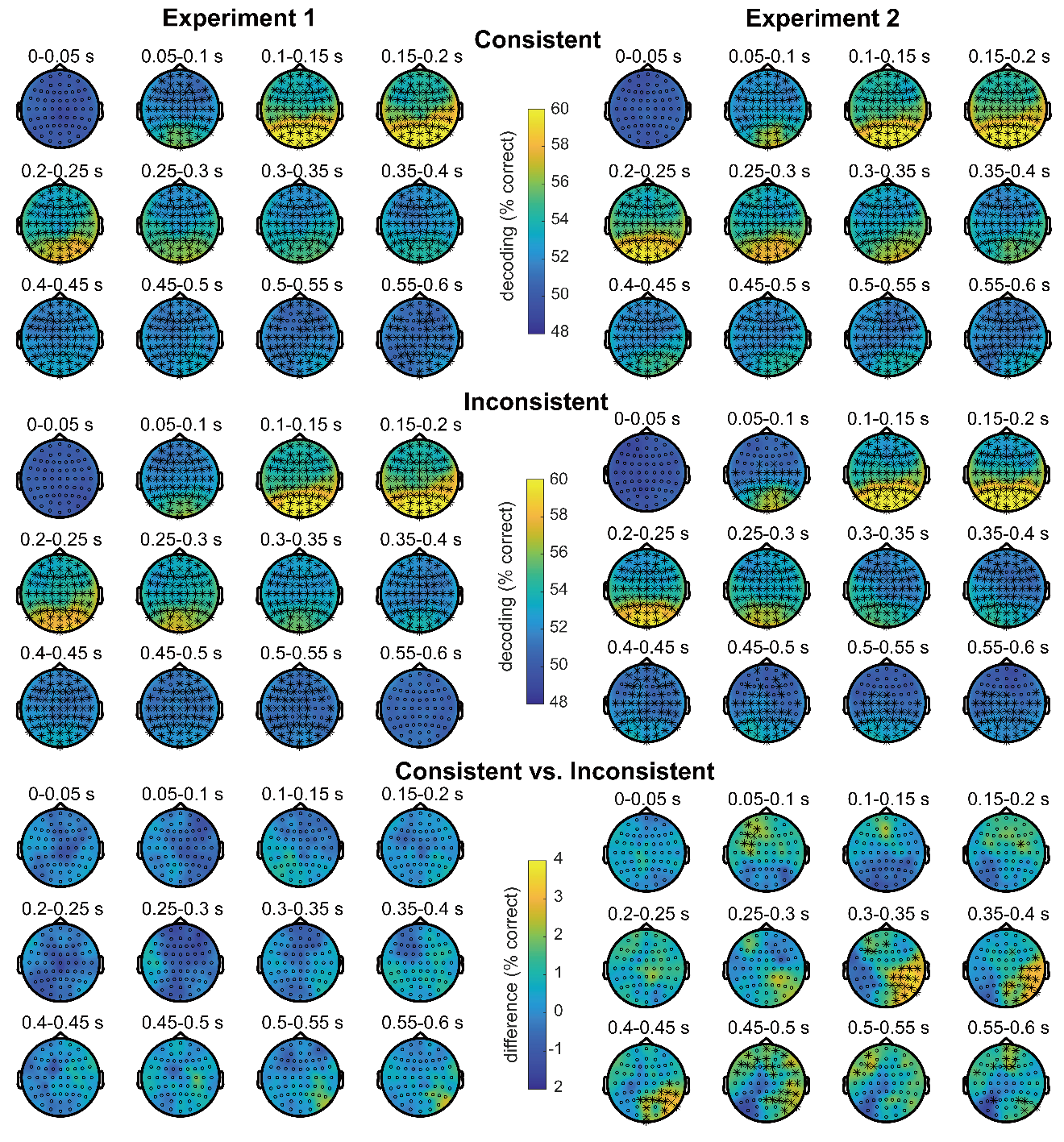

**Fig. S4** Topographies of decoding accuracy for consistent and inconsistent objects in Experiments 1 and 2. Here, the timeseries decoding analysis was repeated for a moving spherical neighborhood of 11 electrodes and accuracies were mapped back onto the scalp location of the central electrode in the neighborhood. The results show that both consistent and inconsistent objects could be decoded across the scalp and from 50-100 ms after object onset, with the strongest decoding in posterior sensors located over visual cortex. Differences between consistent and inconsistent object decoding in Experiment 2 emerged primarily in right-posterior electrodes. For this analysis, electrodes removed during preprocessing were interpolated using data from neighboring electrodes. Asterisks (*) indicate the significant results of comparisons against chance level or between conditions, after FDR correction across electrodes.

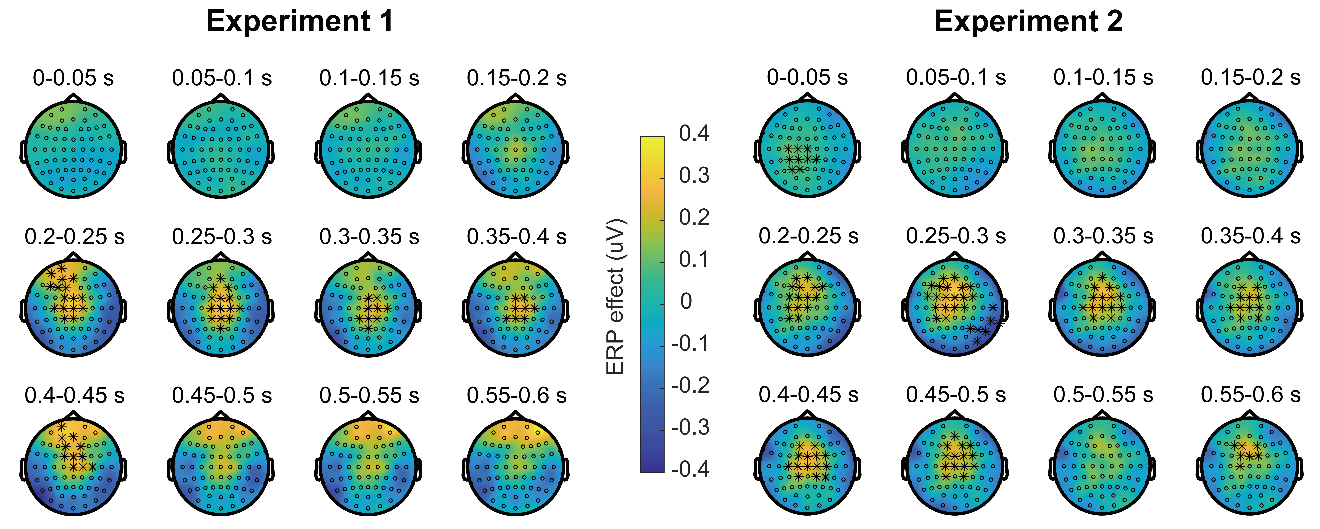

**Fig. S5** Topographies of ERP differences between consistent and inconsistent scene-object combinations in Experiments 1 and 2. Corroborating our electrode selection for main ERP analysis, the most significant scene consistency effects emerged in mid-central region. For this analysis, electrodes removed during preprocessing were interpolated using data from neighboring electrodes. Asterisks (*) indicate the significant differences between the consistent and inconsistent conditions, after FDR correction across electrodes.

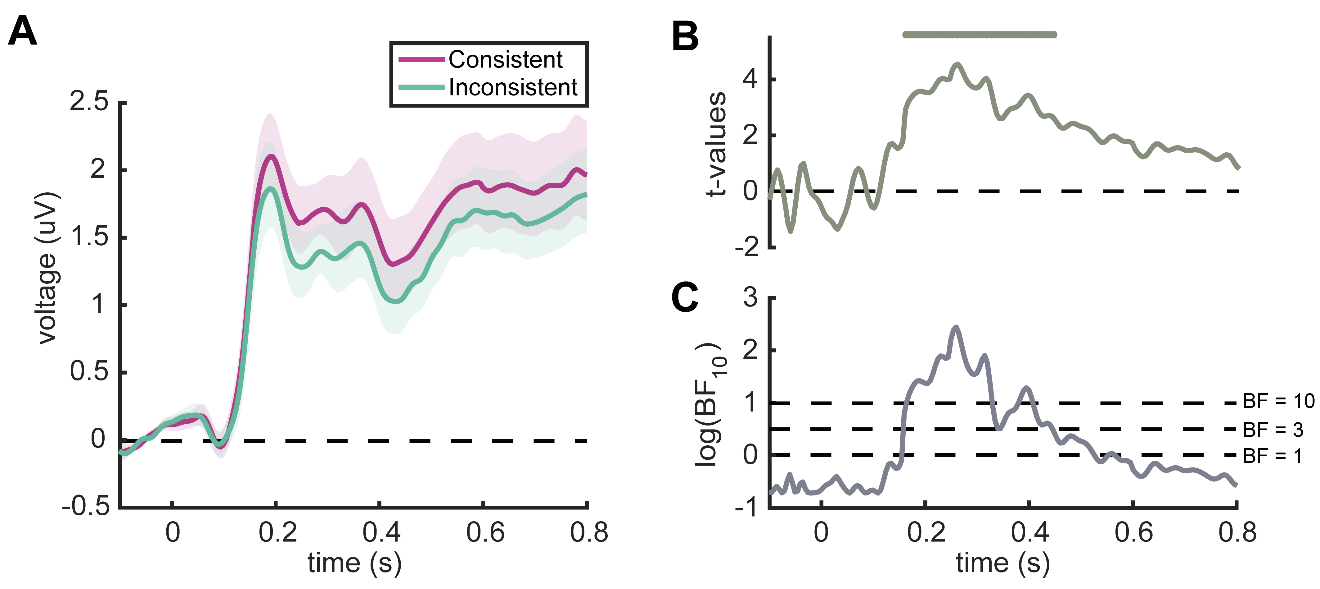

**Fig. S6** Event-related potentials (ERPs) in Experiment 1, with filtering performed before epoching. **(A)** ERPs recorded from the mid-central region for consistent and inconsistent scene-object combinations. Error margins represent standard errors. **(B)** *t*-values for the comparisons between consistent and inconsistent conditions. Line markers denote significant differences between conditions (*p* < 0.05, FDR-corrected). **(C)** Bayes factors (BF_10_) for the comparisons between consistent and inconsistent conditions. For display purposes, the BF_10_ values were log-transformed. Dotted lines show low (BF_10_ = 1), moderate (BF_10_ = 3), and high (BF_10_ = 10) evidence for a difference between conditions. Similar to our original analysis, in which filtering was performed after the other preprocessing steps, inconsistent scene-object combinations evoked more negative responses than consistent combinations at 160-445 ms after object onset.

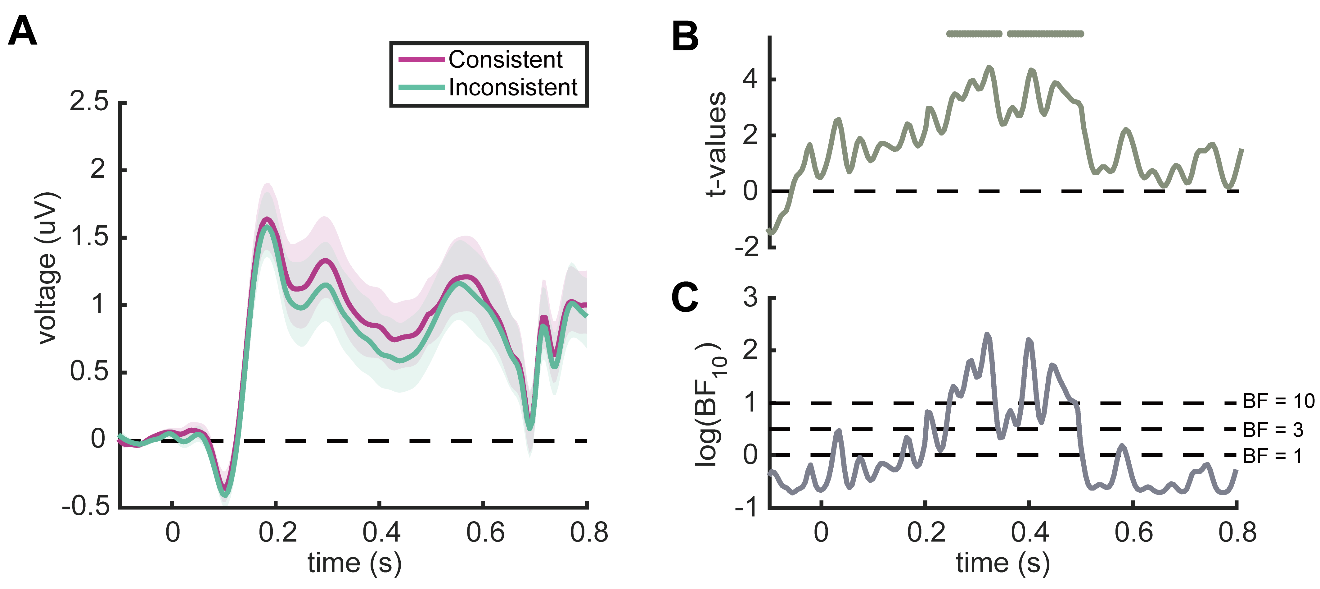

**Fig. S7** Event-related potentials (ERPs) in Experiment 2, with filtering performed before epoching. **(A)** ERPs recorded from the mid-central region for consistent and inconsistent scene-object combinations. Error margins represent standard errors. **(B)** *t*-values for the comparisons between consistent and inconsistent conditions. Line markers denote significant differences between conditions (*p* < 0.05, FDR-corrected). **(C)** Bayes factors (BF_10_) for the comparisons between consistent and inconsistent conditions. For display purposes, the BF_10_ values were log-transformed. Dotted lines show low (BF_10_ = 1), moderate (BF_10_ = 3), and high (BF_10_ = 10) evidence for a difference between conditions. As for Experiment 1, and similar to our original analysis, inconsistent scene-object combinations evoked more negative responses at 245-340 ms and 360-495 ms after object onset, relative to consistent combinations.
